# Meningeal P2X7 signaling mediates migraine-related intracranial mechanical hypersensitivity

**DOI:** 10.1101/2023.02.02.526853

**Authors:** Jun Zhao, Samantha Harrison, Dan Levy

**Author notes:** **Declaration of Interests:** There are no sources of conflict of interest. Author Contributions: Research design: JZ, DL; Conducted experiments: JZ, DL; Performed data analysis: JZ, DL, SH; Writing of manuscript: JZ, DL.

## Abstract

Cortical spreading depolarization (CSD) is a key pathophysiological event that underlies visual and sensory auras in migraine. CSD is also thought to drive the headache phase in migraine by promoting the activation and mechanical sensitization of trigeminal primary afferent nociceptive neurons that innervate the cranial meninges. The factors underlying meningeal nociception in the wake of CSD remain poorly understood but potentially involve the parenchymal release of algesic mediators and damage-associated molecular patterns, particularly ATP. Here, we explored the role of ATP-P2X purinergic receptor signaling in mediating CSD-evoked meningeal afferent activation and mechanical sensitization. Male rats were subjected to a single CSD episode. In vivo, extracellular single-unit recording was used to measure meningeal afferent ongoing activity changes. Quantitative mechanical stimuli using a servomotor force-controlled stimulator assessed changes in the afferent’s mechanosensitivity. Manipulation of meningeal P2X receptors was achieved via local administration of pharmacological agents. Broad-spectrum P2X receptor inhibition, selective blockade of the P2X7 receptor, and its related pannexin 1 channel suppressed CSD-evoked afferent mechanical sensitization but did not affect the accompanying afferent activation response. Surprisingly, inhibition of the pronociceptive P2X2/3 receptor did not affect the activation or sensitization of meningeal afferents post-CSD. P2X7 signaling underlying afferent mechanosensitization was localized to the meninges and did not affect CSD susceptibility. We propose that meningeal P2X7 and Pannexin 1 signaling, potentially in meningeal macrophages or neutrophils, mediates the mechanical sensitization of meningeal afferents, which contributes to migraine pain by exacerbating the headache during normally innocuous physical activities.

**Significance Statement:** Activation and sensitization of meningeal afferents play a key role in migraine headache, but the underlying mechanisms remain unclear. Here, using a rat model of migraine with aura involving cortical spreading depolarization (CSD), we demonstrate that meningeal purinergic P2X7 signaling and its related Pannexin 1 pore, but not nociceptive P2X2/3 receptors, mediate prolonged meningeal afferent sensitization. Additionally, we show that meningeal P2X signaling does not contribute to the increased afferent ongoing activity in the wake of CSD. Our finding points to meningeal P2X7 signaling as a critical mechanism underlying meningeal nociception in migraine, the presence of distinct mechanisms underlying the activation and sensitization of meningeal afferents in migraine, and highlight the need to target both processes for effective migraine therapy.

## Introduction

Migraine is a complex pain syndrome, one of the most common neurological disorders and the leading cause of disability in people under 50 (Collaborators, 2018; Steiner et al., 2018). Migraine consists of episodic attacks characterized by a moderate-to-severe throbbing headache aggravated by physical activity and several other non-painful symptoms, including nausea and light sensitivity. The pathophysiological origin of migraine attacks remains poorly understood, although the headache phase is thought to involve output from trigeminal primary afferent neurons that innervate the cranial meninges (Levy et al., 2019). Persistent discharge of meningeal afferents has been suggested to drive the ongoing migraine headache, while their augmented mechanosensitivity (i.e., sensitization) contributes to migraine pain by exacerbating the headache during normally innocuous physical activities that increase intracranial pressure and drive meningeal mechanical deformations (Gao and Drew, 2016; Blaeser et al., 2022). How meningeal afferents become affected in migraine remain incompletely understood. However, an acute meningeal inflammatory mechanism is considered to play a significant role (Levy et al., 2019).

Cortical spreading depolarization (CSD) is a key pathophysiological event that underlies visual and sensory auras in migraine, which often precede the headache phase (Pietrobon and Moskowitz, 2013). CSD is a self-propagating slow wave of neuronal and glial depolarizations that drastically disrupts transmembrane gradients and cortical synaptic activity (Pietrobon and Moskowitz, 2014). CSD triggering in rodent models produces prolonged activation and mechanical sensitization of meningeal nociceptive afferents (Zhang et al., 2010; Zhao and Levy, 2015, 2016). It is thus an important animal model that can be used to gain insights into the endogenous nociceptive processes that drive meningeal afferents in migraine. In the wake of CSD, numerous algesic mediators are released into the cortical interstitium, including ATP, K^+^, H^+^, arachidonic acid, glutamate, and IL-1β (Csiba et al., 1985; Lauritzen et al., 1990; Schock et al., 2007; Enger et al., 2015; Takizawa et al., 2016; Parker et al., 2021). It has been proposed that these mediators reach the meninges via bulk diffusion or CSF flow, where they can act directly on meningeal afferents or influence their responses indirectly by promoting a meningeal inflammatory response (Carneiro-Nascimento and Levy, 2022).

Here, we focused on ATP signaling as a potential driver of meningeal nociception in the wake of CSD. ATP is a critical algesic molecule that can act directly on nociceptive afferents by activating neuronal P2X2/3 receptors (Staikopoulos et al., 2007; Burnstock, 2016). ATP also serves as a danger-signaling molecule, part of the damage-associated molecular patterns released from stressed or damaged tissues (Di Virgilio et al., 2020), such as the cortex in response to CSD. ATP released in the context of tissue stress occurs partly via the opening of pannexin-1 (Panx1) channels (Dahl et al., 2013), and CSD has been shown to stimulate the opening of Panx1 channels (Karatas et al., 2013). In the cortex, stimulation of P2X7 receptors by ATP controls CSD susceptibility (Chen et al., 2017). However, diffusion of ATP across the cranial meninges and the stimulation of P2X7 expressed by meningeal immune cells (Van Hove et al., 2019) may also drive an inflammatory response and augment the responses of meningeal afferents (McDonald et al., 2010; Roth et al., 2014).

Here, we tested the relative contribution of P2X receptors and Panx1 signaling to the activation and mechanical sensitization of meningeal afferents in a rat model migraine with aura evoked by a single CSD event. Using in vivo extracellular single-unit recording from the trigeminal ganglion in combination with quantitative mechanical stimuli and a pharmacological approach in a rat CSD model, we show that inhibition of the P2X7 receptor and its related Panx 1 channel pore suppresses CSD-evoked meningeal afferent mechanical sensitization. In contrast, local blockade of the pronociceptive P2X2/3 receptor neither affects the activation nor the sensitization of meningeal afferents post-CSD. Additionally, we demonstrate that P2X and Panx1 signaling do not mediate the CSD-evoked acute and persistent discharge of meningeal afferents. Finally, we provide evidence linking meningeal, rather than cortical, P2X7 signaling to meningeal afferent sensitization post-CSD.

## Methods

### Animals

All experiments were approved by the Institutional Animal Care and Use Committee of the Beth Israel Deaconess Medical Centre and followed the ARRIVE (Animal Research: Reporting of In vivo Experiments) guidelines (Kilkenny et al., 2012). Animals (Sprague-Dawley male rats, 220–250 g, Taconic, USA) were housed in pairs with food and water *ad libitum* under a constant 12-hour light/dark cycle (lights on at 7:00 am) at room temperature. All procedures were conducted during the light phase of the cycle (9:00 am to 4:00 pm). Experimental animals were randomly assigned to different treatment groups. No statistical methods were used to predetermine sample size for the experiments, but our sample sizes are similar to those reported in our previous publications using similar techniques (Zhao and Levy, 2016, 2018b; Zhao et al., 2021).

### Surgical preparation for electrophysiological experiments

Animals were deeply anesthetized with urethane (1.5 g/kg, i.p.) and mounted on a stereotaxic frame (Kopf Instruments). A homoeothermic control system kept the core temperature at 37.5–38°C. Animals were intubated and breathed O_2_-enriched room air spontaneously. Physiological parameters were collected throughout the experiments using PhysioSuite (Kent Scientific) and CapStar-100 (CWE). Data used in this report were obtained from animals exhibiting physiological levels of oxygen saturation (> 95%), heart rate (350–450 beats/min), and end-tidal CO_2_ (3.5–4.5%). Two separate craniotomies were made. One was used to expose the left transverse sinus and the posterior part of the superior sagittal sinus, including the adjacent cranial dura, extending ∼2 mm rostral to the transverse sinus. Another small craniotomy (1 × 1 mm) was made over the right hemisphere, centered 2 mm caudal and 2 mm lateral to Bregma, to allow insertion of the recording electrode into the left trigeminal ganglion. A small burr hole (diameter, 0.5 mm) was drilled above the frontal cortex to induce CSD (Zhao and Levy, 2018a). The exposed dura was bathed with a modified synthetic interstitial fluid (SIF) containing 135 mM NaCl, 5 mM KCl, 1 mM MgCl_2_, 5 mM CaCl_2_, 10 mM glucose, and 10 mM HEPES, pH 7.2. In all experiments, SIF was used as the vehicle.

### In vivo recording of meningeal afferent activity

Single-unit activity of meningeal afferents (1 afferent/rat) was recorded from their trigeminal ganglion somata using the contralateral approach reported earlier (Zhao and Levy, 2018b). Meningeal afferents were identified by their constant response latency to electrical stimuli applied to the dura above the ipsilateral transverse sinus (0.5 ms pulse, 1-3 mA, 0.5 Hz). The response latency was used to calculate conduction velocity (CV) based on a conduction distance to the trigeminal ganglion of 12.5 mm (Strassman et al., 1996). Neurons were classified as Aδ (1.5 ≤ CV ≤ 5 m/s) or C afferents (CV < 1.5 m/s). Neural activity was digitized and sampled at 10 kHz using power 1401/Spike 2 interface (CED). A real-time waveform discriminator (Spike 2 software, CED) was used to create a template for the action potential evoked by electrical stimulation to acquire and analyze afferent activity.

### Assessment of meningeal afferent mechanosensitivity

Mechanical receptive fields (RF) of meningeal afferents were first identified using von Frey monofilaments (Zhao and Levy, 2018b). To quantify changes in mechanical responsiveness, we used a servo force-controlled mechanical stimulator (Series 300B Dual Mode Servo System, Aurora Scientific). Mechanical stimuli were delivered using a flat-ended cylindrical plastic probe attached to the tip of the stimulator arm (Zhao and Levy, 2018b). Two ramp-and-hold mechanical stimuli were applied in each trial (rise time, 100 msec; stimulus width, 2 sec; interstimulus interval, 120 sec) and included an initial threshold stimulus (which typically evoked at baseline 1–4 Hz responses) followed by a suprathreshold stimulus (2-3X of the threshold pressure). Stimulus trials were delivered every 15 min throughout the experiment to minimize desensitization (Zhao and Levy, 2018b). Ongoing activity was recorded continuously between the stimulation trials. Basal responses to mechanical stimuli were determined during at least four consecutive trials before the induction of CSD.

### Induction and CSD monitoring

A single CSD episode was induced in the left frontal cortex by briefly inserting a glass micropipette (50 μm diameter) ∼2 mm deep into the cortex for 2 seconds (Zhao and Levy, 2018b). Successful elicitation of CSD was evaluated by simultaneously recording changes in cerebral blood flow with a laser Doppler flowmetry probe positioned within the craniotomy, just above the exposed dura, ∼1 mm from the RF of the recorded unit. The laser Doppler signal was digitized (100 Hz) and recorded using the power 1401/Spike 2. Baseline LDF data were based on recordings conducted for at least 30 min before CSD. Induction of CSD was considered successful when the typical hemodynamic signature characterized by a large transient (∼1 to 2 min) cortical hyperemia, followed by persistent (>1 hour) post-CSD oligemia, was observed (**Figure 1A**) (Gariepy et al., 2017). We investigated drug effect on CSD propagation by measuring the lag time between the pinprick triggering event and the time of maximum cortical hyperemic response associated with the CSD.

**Figure 1.**
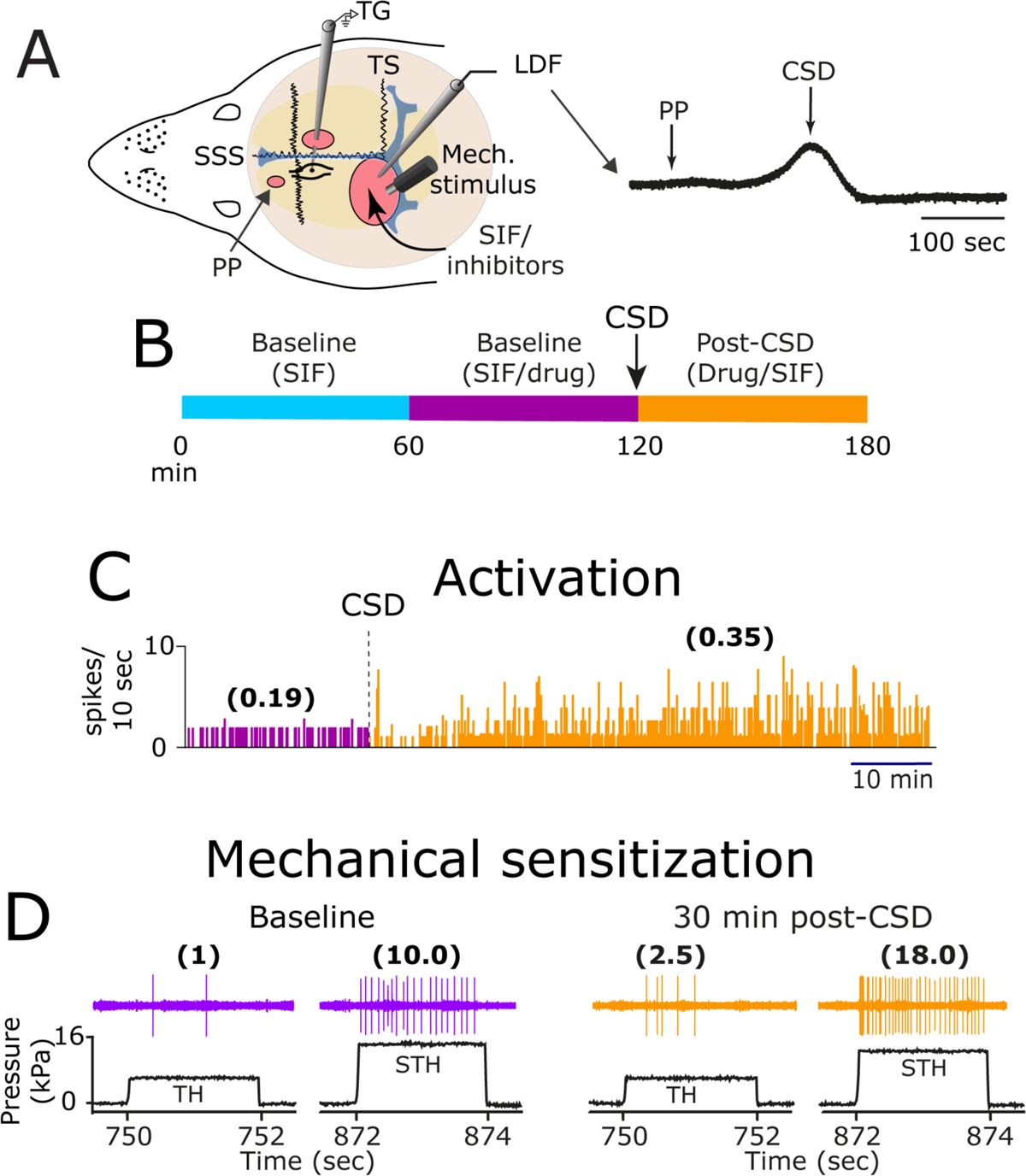
Experimental setup and testing paradigm. **A.** Experimental setup: three skull openings (red ovals) were made. A small burr hole was drilled over the left frontal cortex to elicit a single CSD event using a pinprick (PP) stimulation. Meningeal afferent activity was recorded in the left trigeminal ganglion (TG) using a tungsten microelectrode inserted through a craniotomy over the contralateral hemisphere. An ipsilateral craniotomy was made to expose a part of the left transverse sinus (TS) and superior sagittal sinus (SSS) and their vicinity to find and stimulate mechanosensitive meningeal afferents. Quantitative mechanical stimuli were delivered to the receptive field of afferents using a feedback-controlled mechanical stimulator. A Laser Doppler Flowmeter (LDF) probe was placed over the cortex near the receptive field of afferents to record changes in CBF and validate CSD induction noninvasively. Induction of CSD was considered successful when the typical hemodynamic signature characterized by a large transient (∼1-2 min) cortical cerebral hyperemia, followed by post-CSD oligemia (black trace), was observed. **B.** Experimental paradigm to test the effects of pharmacological blockers. **C.** Examples of an experiment depicting baseline ongoing activity (purple) and the acute and prolonged discharge of meningeal afferents post-CSD (orange) while the meninges are bathed with SIF. Afferent discharge rates in spikes/sec are in parentheses. **D.** Examples of experimental trials depicting the responses of a meningeal afferent to threshold (TH) and suprathreshold (STH) mechanical stimuli (black traces) during baseline recording and then at 30 min after CSD elicitation while the meninges are bathed with SIF. Responses in spikes/sec are in parentheses. Note the CSD-evoked TH and STH sensitization.

### Drugs and treatment approach

We used topical drug application to target local purinergic receptor signaling at the afferents’ meningeal RF level. Drug doses were based on previous studies using local administration and pilot studies in our laboratory. We used Pyridoxalphosphate-6-azophenyl-2’,5’-disulfonic acid tetrasodium salt (PPDAS) as a non-selective P2X antagonist, and 2’,3’-*O*-(2,4,6-Trinitrophenyl)adenosine-5’-triphosphate tetra(triethylammonium) salt (TNP-ATP) to inhibit P2X_3_ and P2X_2,3_ receptors. *N*-[2-[[2-[(2-Hydroxyethyl)amino]ethyl]amino]-5-quinolinyl]-2-tricyclo[3.3.1.13,7]dec-1-ylacetamide dihydrochloride (AZ 10606120) was used to inhibit P2X7, and (3β,20β)-3-(3-Carboxy-1-oxopropoxy)-11-oxoolean-12-en-29-oic acid disodium (carbenoxolone) was employed to inhibit Panx1. All pharmacological agents were purchased from Tocris and diluted in SIF for meningeal application. We first established the level of baseline ongoing activity and mechanosensitivity for 60 min in the presence of the vehicle (SIF). We then evaluated the change in the activity and mechanosensitivity of the afferents in the presence of the drugs for 60 min to ensure a lack of direct inhibitory effect on the afferents before CSD triggering. Post-CSD data was then collected in the presence of the drug or SIF (**Figure 1B**).

### Data and Statistical Analysis

Criteria used to consider meningeal afferent activation and sensitization responses were based on our previous studies (Zhao and Levy, 2018a, b; Zhao et al., 2021). In brief, we considered acute afferent activation when the discharge rate increased ≥ 30 s following the pinprick and lasted ≤ 120 s. A prolonged increase in the ongoing activity of afferents was considered if the firing rate rose above the upper-end point of the 95% CI calculated for the baseline mean for at least 10 consecutive 1 min bins during the first 60 min post-CSD. The development of mechanical sensitization and its duration were based on the following criteria: threshold and/or suprathreshold responses increased to a level greater than the upper endpoint of the 95% CI calculated for the baseline mean; increased responsiveness began during the first 60 min post-CSD, and lasted for at least 30 min.

For analysis of afferent activity, N refers to the number of afferents. Data analyzed using frequentist statistics were plotted using Prism 9 software and are presented in the text as the median and interquartile range (IQR). Group data in graphs show all data points and are plotted as scatter dot plots and the median. Drug effects on CSD-evoked afferent activation and sensitization parameters were analyzed using a two-tailed x^2^ test and non-parametric two-tailed Mann-Whitney or Kruskal-Wallis tests followed by Dunn’s post hoc test. *p* < 0.05 was considered significant.

We employed Bayesian statistics, which is considered to perform better with small sample sizes (McNeish, 2016) as another approach to assess drug effect on CSD-evoked prolonged afferent activation. We used Bayesian hierarchical linear models to compare the effects of vehicle and the different drugs on the CSD-evoked prolonged afferent activation across time (repeated measures for each afferent at 15-60 mins). We adjusted each model for baseline afferent activity, treatment, time, and treatment × time interaction. Afferent responses were considered as a random effect (i.e., random intercept). Prior distributions for all parameters were the default weakly informative values set in the ‘rstanarm’ package for R designed to provide moderate regularization and help stabilize computation. Bayesian estimation was conducted using the default settings with four MCMC chains and 4000 iterations each. The combined posterior distributions were used for inference by extracting the mean and 90% credible interval. We considered treatment effective when a 90% credible interval excluding 0 was present in at least 2 consecutive time points. All analyses and plotting were conducted using RStudio 4.0.

## Results

ATP is a potent pronociceptive mediator that can drive activity in nociceptive afferents, including meningeal afferents (Zhao and Levy, 2015). CSD leads to cortical ATP efflux (Mies and Paschen, 1984; Schock et al., 2007). Astrocytes are a key source of ATP (Xiong et al., 2018), and blocking cortical astrocytic signaling inhibits the development of mechanical sensitization of meningeal afferents post-CSD (Zhao et al., 2021). Hence, we first examined the relative contribution of ATP signaling to the augmentation of meningeal afferent mechanosensitivity post-CSD.

### Broad-spectrum P2X antagonism suppresses CSD-evoked meningeal afferent mechanical sensitization

To begin examining the role of ATP-mediated purinergic signaling, we first tested the effect of local treatment with PPDAS (0.1mM). This broad-spectrum purinergic P2X receptor antagonist has been shown to block the nociceptive effects of ATP (Rong and Burnstock, 2004). PPADS application did not affect CSD triggering (11/11 events), indicating no direct or indirect effects that could influence the vulnerability for CSD induction. We compared CSD-related changes in the mechanical responsiveness of 5 Aδ and 6 C afferents recorded from PPADS-treated animals with 9 Aδ and 13 C afferents recorded from vehicle-treated animals. The data obtained from the Aδ and C afferents were combined as they show similar sensitization characteristics after CSD (Zhao and Levy, 2016). While the afferents’ baseline mechanosensitivity was not affected by PPADS (*p* = 0.62 and *p* = 0.88 vs. control for TH and STH stimuli levels, respectively; Mann-Whitney test; **Table 1**), the post-CSD sensitization was markedly inhibited (**Figure 2**). When mechanical sensitization was defined as enhanced responsiveness at either the TH or STH stimuli levels, we detected fewer sensitized afferents in the PPADS-treated group compared to the control group (2/11 vs. 17/23 sensitized afferents; χ^2^ = 9.3, *p* = 0.002, **Figure 2B**]. PPADS treatment also inhibited afferent sensitization, defined as enhanced responsiveness at both TH and STH stimuli levels (PPADS: 0/11, Control: 10/23 sensitized afferents; χ^2^ = 6.8, *p* = 0.009, **Figure 2C**). PPADS also decreased the overall changes in TH and STH responses post-CSD when compared to the vehicle [TH, −0.3 (1.6) spikes/sec vs. 0.9(1.7) spikes/sec, *p* = 0.048; STH, 0.9(0.2) vs. 1.3 (0.7) fold, *p* = 0.004, Mann-Whitney test, **Figures 2D, E**].

**Figure 2.**
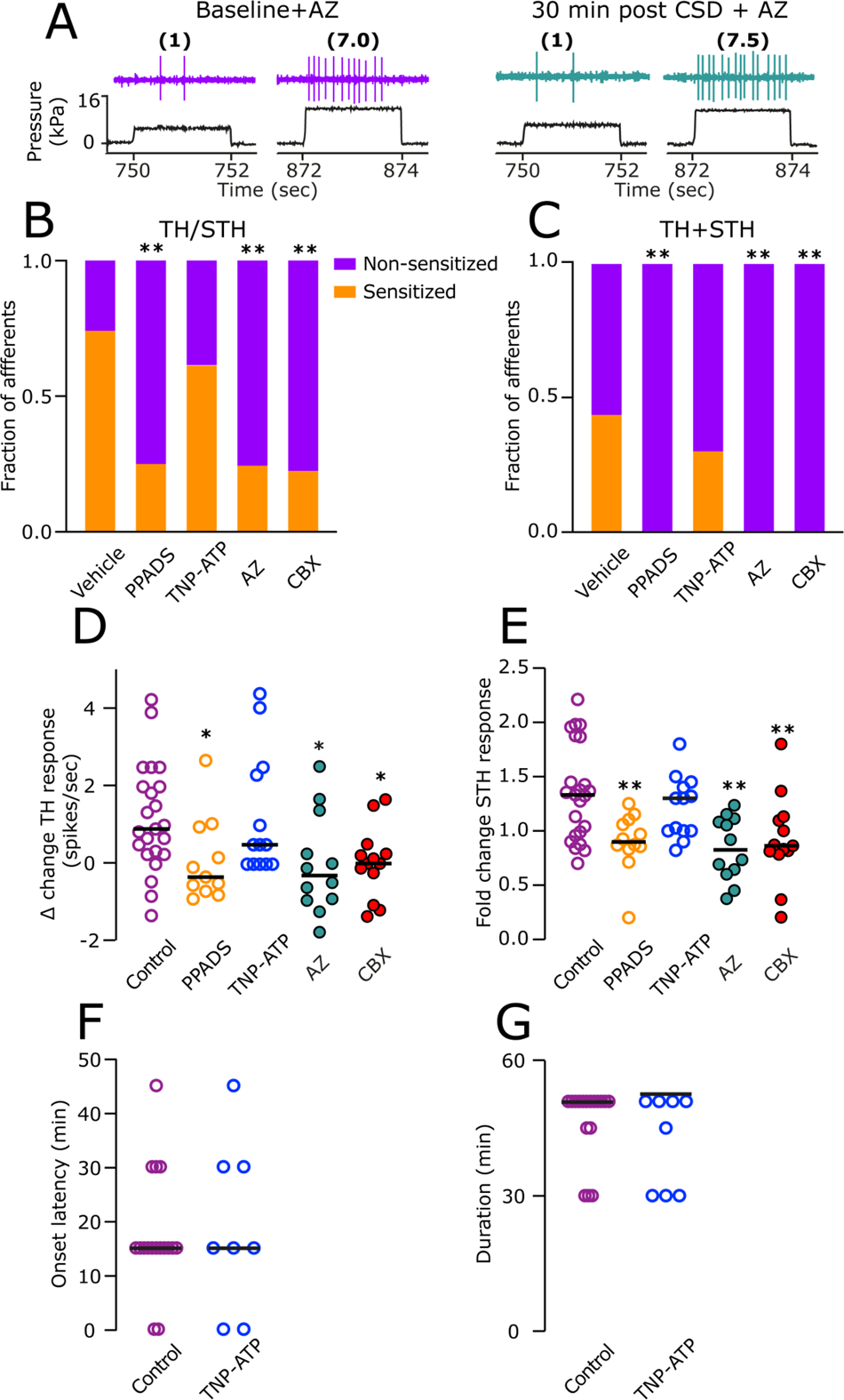
Inhibition of P2X7 receptors and Panx1 channels suppresses CSD-evoked mechanical sensitization of meningeal afferents. **A.** Examples of experimental trials depicting the Threshold (TH) and suprathreshold (STH) mechanical responses of a meningeal afferent during baseline recording and then at 30 min after CSD elicitation in the presence of the P2X7 receptor inhibitor AZ10606120 (AZ). Note the lack of sensitization at the TH and STH stimuli levels. When compared to controls (n=22), the fraction of afferents that became sensitized at either TH or STH levels (TH/STH, **B**) or at both levels (TH+STH, **C**) was lower in animals treated with the broad-spectrum purinergic P2X receptor antagonist PPADS (n=18), AZ10606120 (n=12), or the Panx1 inhibitor, carbenoxolone (CBX, n=13), * *p* < 0.05, χ^2^ test, treatment vs. control). Inhibition of P2X2/3 and P2X3 receptors with TNP-ATP (n=13) did not affect the sensitization response. At the population level, inhibition of P2X, P2X7, or Panx1 also decreased the magnitude of the TH **(D)** and STH **(E)** afferent responsiveness post-CSD. Significance *p* values indicate Dunn’s post hoc test between a given treatment and control following a significant (** *p* < 0.01) Kruskal-Wallis test between all treatments. TNP-ATP treatment did not affect the afferents’ TH and STH response magnitude post-CSD (**D, E** *p* > 0.05). TNP-ATP also did not affect the onset latency **(F**) and duration of the post-CSD sensitization response (**G**) when compared to vehicle control, *p* > 0.05 Mann Whitney test. Data in **D-G** include all data points. Lines represent the median.

**Table 1:** Baseline mechanical response properties of meningeal afferents in animals treated with vehicle (control), PPADS, TNP-ATP, AZ10606120 (AZ), Carbenoxolone (CBX)

|  | **Meningeal**<br>**mechanical RFs** | **TH responses** | **STH responses** |
| --- | --- | --- | --- |
| Vehicle (n=23) | 2(1) | 1.5(2.2) | 8.0(6.5) |
| PPADS (n=11) | 2(2) | 1.8 (1.8) | 7.8(5.6) |
| TNP-ATP (n=13) | 2(1) | 2.0(1.4) | 7.3(11.0) |
| AZ (n=12) | 2(1) | 2.5(1.5) | 7.0(6.1) |
| CBX (n=12) | 2(2) | 2.0(1.6) | 8.3(10.6) |

Data are expressed as median (IQR). N represents the number of afferents tested (1 afferent/animal). The number of distinct dural receptive fields (RFs) was tested using a suprathreshold von Frey stimulus. Threshold (TH) and suprathreshold (STH) responses (in spikes/second) are based on activity recorded during a 2-s stimulation and represent the average of 4 trials conducted before CSD while the afferent RF was bathed with vehicle or drug. Group differences were analyzed using the 2-tailed Kruskal–Wallis test and were not significant ( $p > 0.05$ ) for the 3 parameters tested.

### Inhibition of P2X2/3 does not affect CSD-evoked meningeal afferent mechanical sensitization

We next tested the possibility that ATP produces meningeal afferent sensitization by acting upon P2X2/3, the principal P2X receptors expressed by meningeal and other primary afferent nociceptive neurons (Staikopoulos et al., 2007; Burnstock, 2016). We examine the effect of TNP-ATP (0.1 mM), a potent P2X2/3 receptor antagonist (Jarvis et al., 2001), which has been shown to block ATP-mediated nociceptive neuron responses ((Burgard et al., 1999), including of meningeal afferents, see below). Topical application of TNP-ATP neither affected CSD triggering (13/13 events) nor the afferent sensitization response. Among the afferent tested (6 Aδ; 7 C), when sensitization was defined as enhanced responsiveness at either the TH or STH stimuli level, we observed sensitization in 8/13 afferent, not different than in controls (χ^2^ = 0.6, *p* = 0.44, **Figure 2B**). Similarly, TNP-ATP did not inhibit the CSD-evoked sensitization defined as enhanced responsiveness at both the TH and STH stimuli levels (3/13 sensitized afferents, χ^2^ = 1.5, p = 0.22, **Figure 2C**]. TNP-ATP treatment also resulted in similar sensitization magnitudes [TH, 0.5(2.4) spikes/sec, *p* = 0.99 vs. control; STH, 1.3(0.4) fold, *p* = 0.54 vs. control, Mann Whitney; **Figure 2D, E**], latencies for developing sensitization [(15.0(26.5) min vs. (15 (7.5), *p* = 0.89, and durations of the sensitized state [52.5(30.0) vs. 60(15) min, *p* 0.33, Mann-Whitney test; **Figure 2F, G**].

### Blockade of P2X7 and Panx1 signaling inhibits CSD-evoked meningeal afferent mechanical sensitization

P2X7 receptors are expressed in numerous meningeal immune cells and mediate proinflammatory functions (Van Hove et al., 2019). Because a broad spectrum P2X antagonism (PPADS), but not P2X2,3 inhibition (TNP-ATP), affected CSD-evoked afferents’ mechanosensitization, we next asked whether P2X7 might be involved. To test the contribution of this purinergic receptor, we used the selective P2X7 receptor antagonist AZ10606120 (10 μm), which has been shown to block proinflammatory responses (Sluyter and Vine, 2016). We recorded CSD-evoked responses of 4 Aδ and 8 C afferents in animals treated topically with AZ10606120. We observed no effect on eliciting single CSD events (12/12 events) or changes in the afferents’ baseline mechanosensitivity (**Table 1**). Administration of AZ10606120, however, significantly decreased the afferents’ propensity to develop mechanical sensitization post-CSD, both when sensitization was defined as increases at TH or STH level (3/12 sensitized afferents, χ^2^ = 7.7, *p* = 0.006 vs. control, **Figure 2B**) or at TH plus STH levels (0/12 sensitized afferents, χ^2^ =7.3, *p* = 0.007 vs. control, **Figure 2C**). The post-CSD response magnitudes in animals pretreated with the P2X7 antagonist were also diminished when compared to vehicle-treated controls [TH, −0.3(2.9) spikes/sec, *p* = 0.04 vs. control; STH, 0.8 (0.5) fold, *p* = 0.001 vs. control, Mann-Whitney test; **Figure 2D, E**].

P2X7 forms a complex with Panx1 channels that serve as a large pore channel and conduit for the further release of ATP plus other proinflammatory mediators, such as IL-1β (Mehta et al., 2001). Several meningeal immune cells, including macrophages, dendritic cells, and T cells, express Panx1 together with P2X7 (Van Hove et al., 2019), pointing to its potential contribution to afferent sensitization post-CSD. Because P2X7 antagonists may also inhibit the P2X7/Panx1 pore activity and related ATP release (Xia et al., 2012; Leeson et al., 2018), we next tested the effect of the Panx1 inhibitor carbenoxolone (CBX, 1mM) (Michalski and Kawate, 2016) on the responses of 7 Aδ and 6 C afferents. Topical CBX application did not affect CSD induction (13/13 events) or basal afferent mechanosensitivity (**Table 1**). Treatment with CBX, however, diminished the propensity of the afferents to develop mechanical sensitization when considering changes at the TH or STH levels, 3/13 sensitized afferents, χ^2^ =8.7, *p* = 0.003 vs. control, **Figure 2B**) or at both levels (0/13 sensitized afferents, χ^2^ = 7.8, *p* = 0.005 vs. control, **Figure 2C**). The post-CSD changes in TH and STH responses in CBX-treated animals were also reduced when compared to controls (TH, 0.0(0.9) spikes/sec, *p* = 0.03 vs. control; STH, 0.9 (0.3) fold, *p* = 0.004 vs. control, Mann-Whitney test; **Figure 2D, E**).

### CSD-evoked meningeal afferent activation does not involve P2X receptor signaling or Panx1 activation

The mechanisms underlying meningeal afferent mechanical hyperresponsiveness following CSD appear independent of those responsible for the increased ongoing discharge (Zhao and Levy, 2016, 2018b). We, therefore, asked next whether P2X and/or Panx1 inhibition might also affect the CSD-evoked increases in the afferents’ ongoing activity. We first analyzed the effect of the pharmacological inhibitors on the development of the acute afferent discharge that develops during or immediately after the passage of the CSD wave under the afferents’ meningeal receptive field. Topical application of PPDAS neither affected the afferents’ baseline ongoing activity (**Table 2**) nor their acute activation in response to CSD. We observed acute activation in 3/8 Aδ and 3/7 C afferents, not different from the vehicle control group 7/24 Aδ and 11/32 C; x^2^ = 0.3, *p* = 0.57 for the combined population, **Figure 3B**).

**Figure 3.**
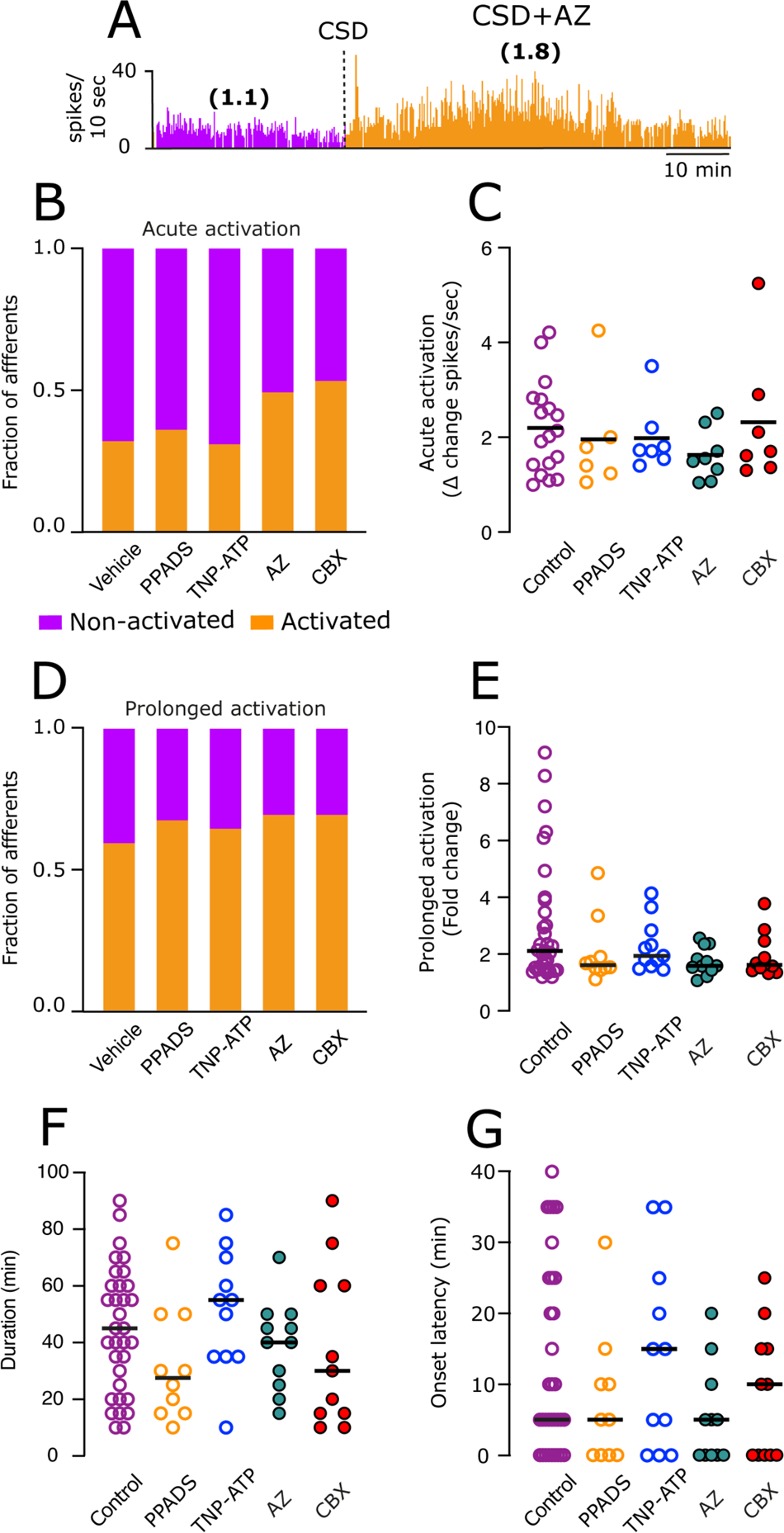
CSD-evoked meningeal afferent activation is not affected by pharmacological inhibition of P2X receptors and Panx1. **A**. Example experimental trial depicting the CSD-evoked acute and prolonged activation of a meningeal afferent in an AZ10606120-treated animal. Ongoing activity (spikes/sec) is in parentheses. **B.** Broad spectrum P2X inhibition, selective antagonists of P2X2/3, P2X7, or Panx1 blockade did not affect the propensity of the afferents to develop acute activation (*p* > 0.05, χ^2^ test for each treatment vs. control). **C.** The magnitude of the acute response was also not affected (*p* > 0.05 for all comparisons, Kruskal-Wallis test). **D.** Inhibition of P2X, P2X2/3, P2X7, or Panx1 also did not affect the propensity of the afferents to develop prolonged activation (*p* > 0.05, χ^2^ test for each treatment vs. control). The magnitudes **(E)**, durations **(F),** and onset latencies **(G)** of the prolonged response were also not affected (*p* > 0.05 Kruskal-Wallis test for all comparisons vs. control). Data in **C, E-G** include all data points. Lines represent the median.

**Table 2:** Baseline ongoing activity in animals treated with vehicle, PPADS, TNP-ATP, AZ10606120, or Carbenoxolone.

|  | **Aδ** | **C** | **Combined population** |
| --- | --- | --- | --- |
| Vehicle | (n=23); 0.1(0.2) | (n=33); 0.6(1.0) | 0.2(0.8) |
| PPADS | (n=8); 0.11 (0.7) | (n=7); 1.1(0.6) | 0.5 (1.1) |
| TNP-ATP | (n=8); 0.1(0.7) | (n=9); 0.7(0.7) | 0.3(0.7) |
| AZ10606120 | (n=7); 0.1 (0.8) | (n=8); 0.6(0.4) | 0.4(0.7) |
| Carbenoxolone | (n=8); 0.1 (0.4) | (n=8); 1.2(1.1) | 0.4 (1.2) |

Data are expressed as median (IQR). N represents the number of afferents tested (1 afferent/animal). Ongoing activity (in spikes/sec) is based on data collected for at least 45 minutes before CSD while the RF of the afferents was bathed with the drug or vehicle (SIF). Group differences were analyzed using 2-tailed Kruskal–Wallis test and were not significant ( $p > 0.05$ ) for the population tested.

We next tested the effect of the P2X2/3 inhibitor TNP-ATP on the CSD-evoke afferent activation. In the presence of TNP-ATP, there was no change in baseline ongoing activity (**Table 2**), and acute CSD-evoked afferent activation was also unaffected (5/8 Aδ and 2/9 C afferents, x^2^ = 0.5 *p* = 0.49 vs. controls for the combined population, **Figure 3B**). While TNP-ATP is considered a potent P2X2/3 inhibitor (Jarvis et al., 2001), to verify that the lack of effect we observed was not due to an insufficient dose, we tested whether local application of 0.1 mM TNP-ATP can inhibit ATP-evoked acute discharge of meningeal afferents. When compared to historical data showing acute afferent activation by ATP (1 mM in the presence of vehicle, 17/30 afferents (Zhao and Levy, 2015)), pretreatment plus co-administration of TNP-ATP (0.1 mM) blocked the excitatory effect produced by local ATP administration at 1mM (2/10 afferents, x^2^ = 4.0, *p* = 0.04 vs. control) pointing to a potent inhibitory effect of TNP-ATP using this concentration on meningeal afferents.

Local application of the selective P2X7 receptor antagonist AZ10606120 also did not inhibit baseline ongoing activity (**Table 2**) or the acute afferent response (3/7 Aδ and 5/9 C-afferents, x^2^ = 1.7 *p* = 0.19 vs. controls). Similarly, Panx1 inhibition with CBX neither affected the baseline ongoing activity (**Table 2**) nor the acute response resulting in the activation of 4/8 Aδ and 3/8 C (x^2^ = 0.7 *p* = 0.39 vs. controls, **Figure 3B**). None of the inhibitors tested affected the magnitude of the acute activation (H_5_ = 2.76; *p* = 0.59, Kruskal-Wallis test, **Figure 3C**).

We next examined whether inhibition of purinergic receptors and Panx1 affect the development of prolonged afferent activation post-CSD. None of the inhibitors affected the prolonged increase in afferent discharge (**Figure 3D).** Following PPDAS treatment, we detected prolonged activation in 5/8 Aδ and 5/7 C afferents, not different from the activation observed in vehicle-treated animals (13/24 Aδ, 20/32 C; x^2^ = 0.3 *p* = 0.59 for the combined population). The propensity to develop prolonged activation post-CSD was also unaffected by TNP-ATP treatment (5/8 Aδ, 6/9 C; x^2^ = 0.2 *p =* 0.67 for the combined population vs. control). Treatment with AZ10606120 resulted in prolonged activation in 4/7 Aδ and 7/9 C-afferents. Also, not different from the control group (x^2^ = 0.5 *p* = 0.48 for the combined population). Finally, CBX application failed to inhibit the prolonged activation, which we observed in 5/8 Aδ and 6/8 C-afferents (x^2^ = 0.5 *p* > 0.48 vs. controls for the combined population). We next analyzed whether P2X or Panx1 inhibition might affect any of the prolonged activation parameters. However, none of the drugs tested affected the prolonged afferent activation magnitude, duration, or onset latency [(H_5_ = 5.4, *p* = 0.25; H_5_, *p* = 0.48; H_5_, *p* = 0.19 respectively, Kruskal-Wallis tests, **Figure 3E, F, G**)].

Because our frequentist statistics approach did not reveal any drug effect on the increased afferent activity following CSD, we also analyzed the data using Bayesian statistics. However, the Bayesian hierarchical linear models also failed to detect differences between the changes in afferent activity observed in the vehicle and any drug treatment groups (**Table 3**).

**Table 3.**
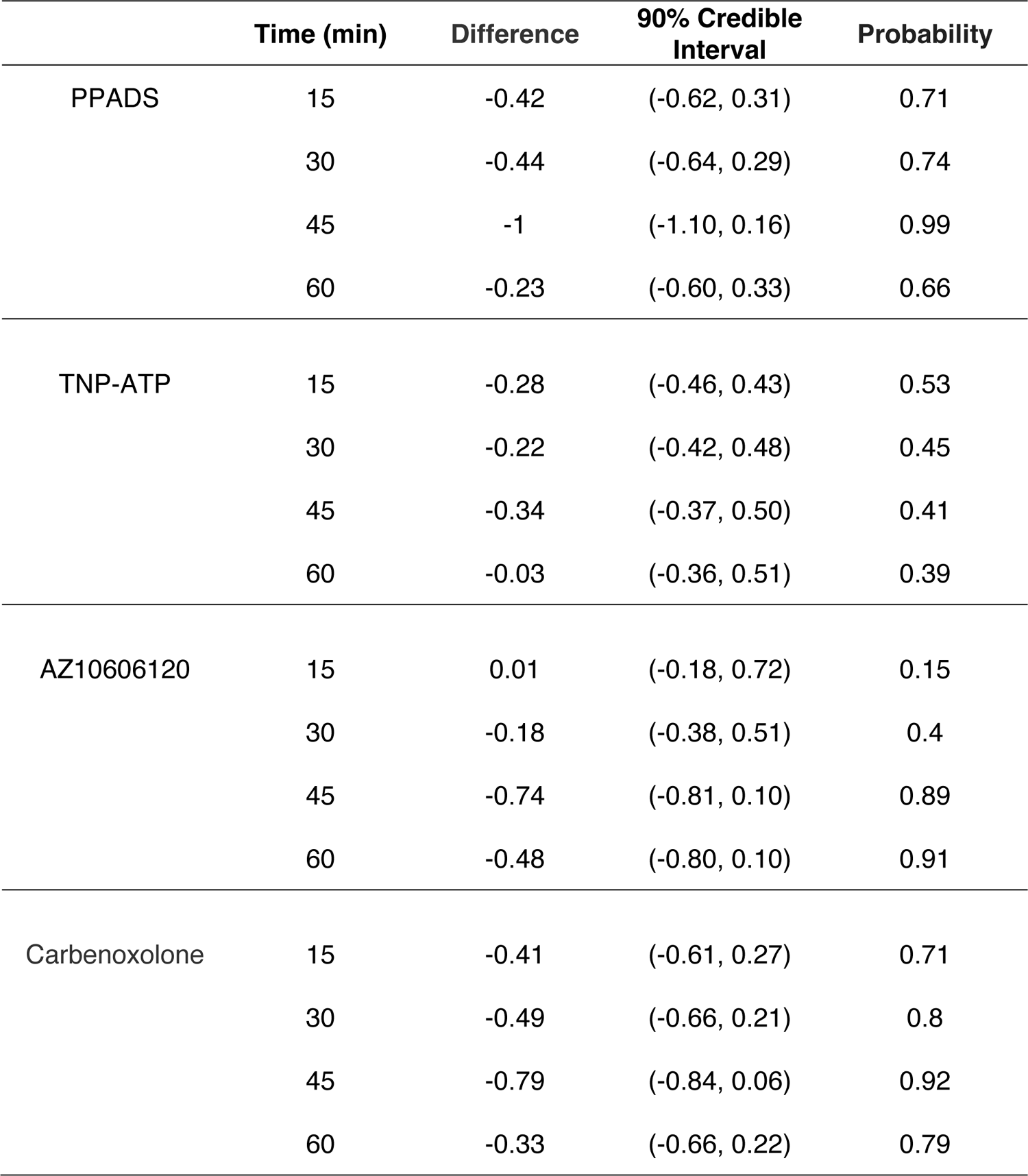
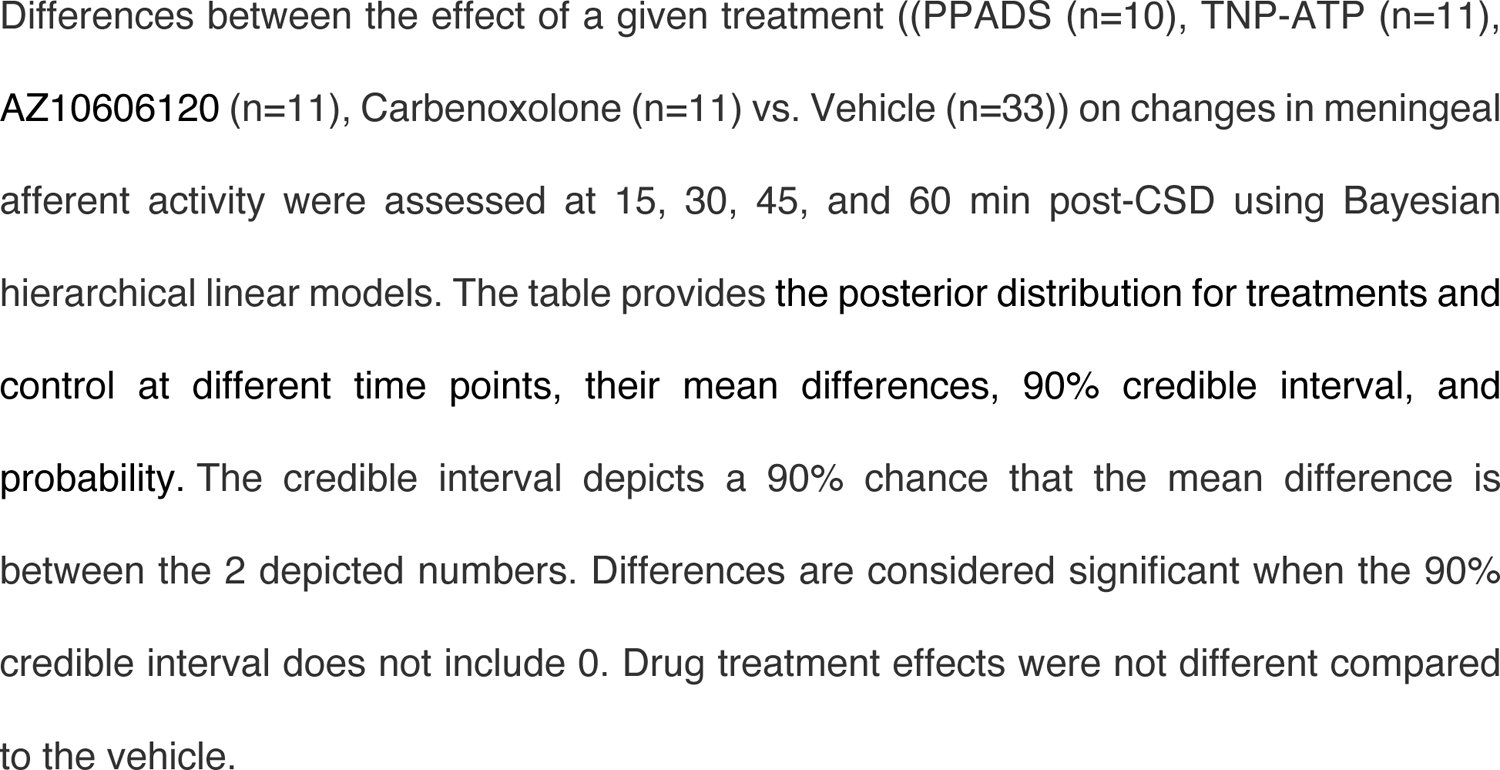
Bayesian posterior probability distributions of models used to compare the effects of PPADS, TNP-ATP, AZ10606120, and Carbenoxolone vs. vehicle on CSD-evoked prolonged meningeal afferent activation.

|  | **Time (min)** | **Difference** | **90% Credible Interval** | **Probability** |
| --- | --- | --- | --- | --- |
| PPADS | 15 | -0.42 | (-0.62, 0.31) | 0.71 |
|  | 30 | -0.44 | (-0.64, 0.29) | 0.74 |
|  | 45 | -1 | (-1.10, 0.16) | 0.99 |
|  | 60 | -0.23 | (-0.60, 0.33) | 0.66 |
| TNP-ATP | 15 | -0.28 | (-0.46, 0.43) | 0.53 |
|  | 30 | -0.22 | (-0.42, 0.48) | 0.45 |
|  | 45 | -0.34 | (-0.37, 0.50) | 0.41 |
|  | 60 | -0.03 | (-0.36, 0.51) | 0.39 |
| AZ10606120 | 15 | 0.01 | (-0.18, 0.72) | 0.15 |
|  | 30 | -0.18 | (-0.38, 0.51) | 0.4 |
|  | 45 | -0.74 | (-0.81, 0.10) | 0.89 |
|  | 60 | -0.48 | (-0.80, 0.10) | 0.91 |
| Carbenoxolone | 15 | -0.41 | (-0.61, 0.27) | 0.71 |
|  | 30 | -0.49 | (-0.66, 0.21) | 0.8 |
|  | 45 | -0.79 | (-0.84, 0.06) | 0.92 |
|  | 60 | -0.33 | (-0.66, 0.22) | 0.79 |

Differences between the effect of a given treatment ((PPADS (n=10), TNP-ATP (n=11), AZ10606120 (n=11), Carbenoxolone (n=11) vs. Vehicle (n=33)) on changes in meningeal afferent activity were assessed at 15, 30, 45, and 60 min post-CSD using Bayesian hierarchical linear models. The table provides the posterior distribution for treatments and control at different time points, their mean differences, 90% credible interval, and probability. The credible interval depicts a 90% chance that the mean difference is between the 2 depicted numbers. Differences are considered significant when the 90% credible interval does not include 0. Drug treatment effects were not different compared to the vehicle.

### The meningeal anti-nociceptive effect mediated by P2X7 inhibition does not involve cortical action

ATP diffused from the cortex can drive the activation of meningeal immune cells expressing P2X7 and indirectly contribute to afferent sensitization. However, activation of cortical astrocytic, microglial, and neuronal P2X7 receptors (Illes et al., 2017) might also play a role. Although we applied the pharmacological inhibitors topically to the meninges, it is technically challenging to ascertain whether they also diffused into the cortex. Agents applied topically to the meninges can diffuse into the CNS. This approach, however, results in a much lower concentration of the agents in the cortex and CSF, with an estimated dilution factor of approximately 10^-5^ (Zhao et al., 2017), which is expected to limit their cortical action. Nonetheless, we conducted an additional experiment to test whether AZ10606120 applied to the dura mater might also affect cortical P2X7-mediated responses.

Previous studies documented suppression of KCl-evoked CSD triggering in response to cortical P2X7/Panx1 pore activity inhibition (Chen et al., 2017) but not after Panx1 inhibition with CBX (Karatas et al., 2013). Building upon this previous data, we examined whether local application of AZ10606120 at the same concentration that inhibited meningeal afferent sensitization might suppress the triggering of multiple CSD events. We observed that AZ10606120 administration (n=7) did not affect the generation of multiple CSDs in response to continuous cortical KCl stimulation when compared to vehicle (n=6) treatment [AZ10606120 11.0 (3.0) CSDs/hr; vehicle 11.5 (3.7) CSDs/hr, *p* = 0.76, Mann Whitney test; **Figure 4A-C**] suggesting that the effect of the topically applied AZ10606120 on the sensitization of meningeal afferents involved a meningeal action.

**Figure 4.**
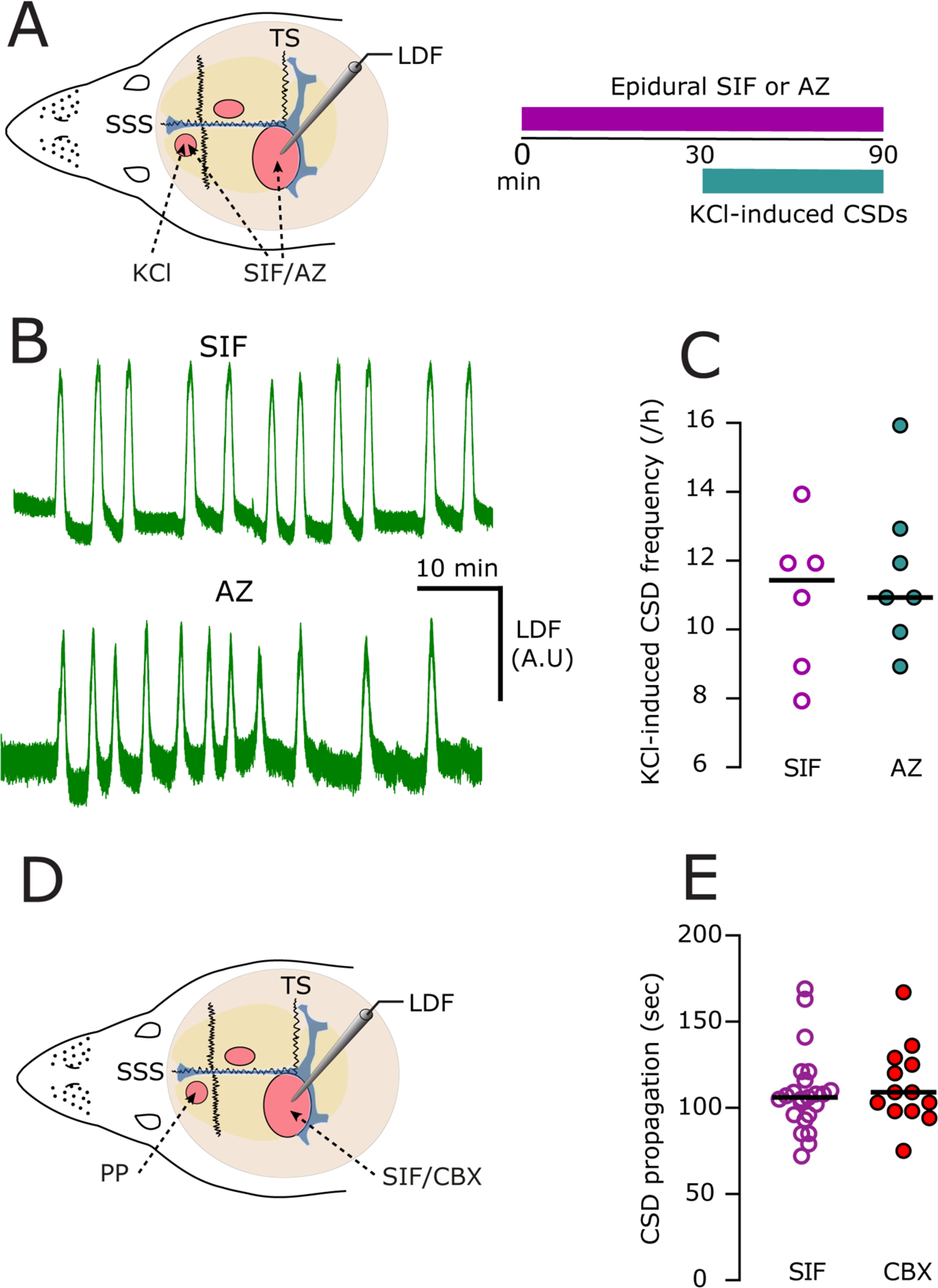
Epidural application of AZ10606120 (AZ) does not suppress the frequency of KCl-evoked cortical spreading depolarization (CSD). (**A**) Experimental setup and testing paradigm. **A.** We used the same surgical approach as in the TG recording, including three skull openings (red ovals). Continuous application of 1M KCl to the anterior burr hole was used to elicit multiple CSD events, monitored using laser Doppler flowmetry (LDF) in the posterior opening. The burr hole and the site of LDF monitoring were pretreated with SIF or the P2X7 inhibitor (AZ10606120, 10 μm) for 30 min. CSD frequency was measured during the application of KCl in SIF or KCl+AZ. **B.** Representative LDF recordings show similar CSD frequencies in animals treated with SIF or AZ10606120. **C.** Scatter dot plot showing that P2X7 inhibition with AZ10606120 (n=7) does not inhibit KCl-evoked spreading depression frequency compared to vehicle (n=6). Data include all data points. Lines represent the median. *p* > 0.05 Mann-Whitney test.

In addition to blocking Panx1, CBX also inhibits gap junctions (Spray et al., 2019), which potentially could also impact the CSD-evoked meningeal nociceptive effect we observed. Although cortical action of CBX does not inhibit CSD triggering (Karatas et al., 2013), its impact on gap junctions affects CSD propagation velocity (Theis et al., 2003; Tamura et al., 2011). Hence, to explore whether CBX diffusion following topical meningeal application might influence cortical gap junctions, we examined its effect on the propagation of CSD from the rostral triggering site to the recording site (**Figure 5A**). When compared to vehicle treatment (n=23), CBX (n=13) did not affect CSD propagation [CBX 109.0 (29.0) sec; vehicle 106.0 (20.0) sec, *p* = 0.55, Mann-Whitney test; **Figure 5B**], suggesting that topically administered CBX may not inhibit the sensitization of meningeal afferent post-CSD by blocking cortical gap junctions.

**Figure 5.**
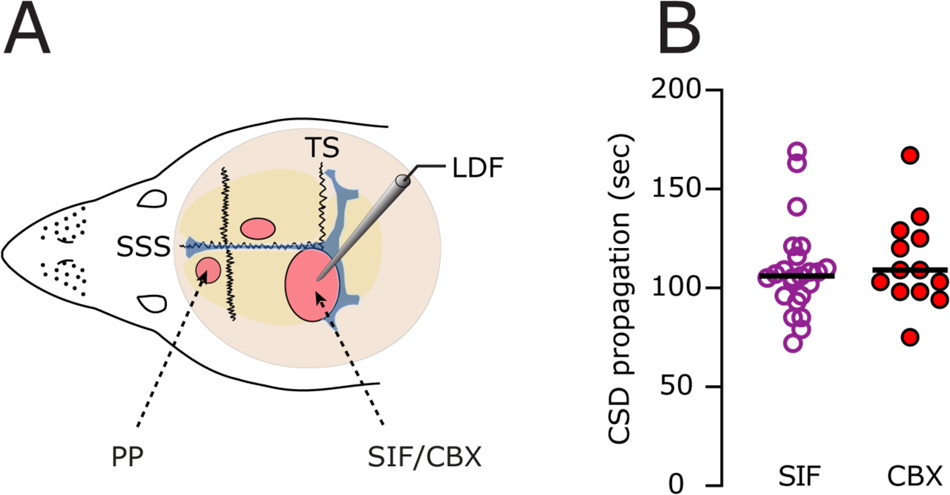
Epidural application of Carbenoxolone (CBX) does not affect CSD propagation. **A.** Experimental approach to test the effect of topical meningeal CBX treatment on CSD propagation from the rostral pinprick (PP) triggering site to the caudal recording site, where CSD-related cortical hyperemia was measured with laser Doppler flowmetry (LDF). TS - transverse sinus; SSS - superior sagittal sinus. **B.** Scatter dot plot showing that tropical application of CBX (1 mM, n=13) does affect CSD propagation compared to vehicle (n=23), suggesting no effect on cortical gap junctions. Data include all data points. Lines represent the median. *p* > 0.05 Mann-Whitney test.

## Discussion

The headache phase in migraine with aura involves the activation and mechanical sensitization of meningeal afferents in response to a CSD event. However, the mechanisms by which CSD influences the responses of meningeal afferents have yet to be fully elucidated. Here we show that CSD augments the mechanosensitivity of meningeal afferents via the purinergic P2X7 channel and its related Panx1 pore. We further provide evidence that this meningeal anti-nociceptive effect is mediated by meningeal and not cortical P2X7-related signaling.

Our findings that blocking P2X7 and Pannexin 1 interfere with the development of mechanical sensitization but does not impede the increase in afferent discharge post-CSD agrees with our previous data showing that the activation and sensitization of meningeal afferents, including in response to CSD, involves distinct mechanisms (Zhao and Levy, 2016, 2018b; Zhao et al., 2021).

The mechanism by which P2X7/Panx1 activation drives meningeal afferent sensitization remains to be explored but likely involves an inflammatory response. Single-cell RNA sequencing data demonstrated the expression of P2X7 plus Panx1 in meningeal macrophages (Van Hove et al., 2019), pointing to these immune cells as critical factors mediating afferent sensitization post-CSD. P2X7 and Panx1 signaling were also implicated in meningeal neutrophil recruitment under pathological conditions (Roth et al., 2014) and thus could mediate the meningeal nociceptive response produced by CSD. Meningeal P2X7/Panx1 activation can lead to the release of the proinflammatory and pronociceptive cytokines-1β and TNF-α plus the activation of phospholipase A2 and the release of sensitizing prostanoids (Mariathasan et al., 2006; Norris et al., 2014; Barbera-Cremades et al., 2017). In turn, IL-1β, TNF-α, and prostanoids sensitize meningeal afferents (Zhang et al., 2011; Zhang et al., 2012; Zhao and Levy, 2018b).

CSD gives rise to persistent vasodilatation of the middle meningeal artery, a process proposed to be mediated by prolonged activation of peptidergic meningeal afferents (Bolay et al., 2002). Karatas and colleagues (Karatas et al., 2013) have shown that the local blockade of Panx1 with CBX suppresses this meningeal vasodilatory response. Our finding that CBX did not inhibit the CSD-evoked prolonged meningeal afferent activation argues, however, for a dissociation between the CSD-evoked dural vasodilatation enhanced ongoing meningeal afferent discharge. Whether signaling linked to the increased afferent mechanosensitivity post-CSD also exerts a local vasodilatory effect requires further investigation.

ATP is an algesic mediator that excites nociceptors, including meningeal afferents, by activating neuronal purinergic P2X2/3 receptors. Surprisingly, blocking this receptor using a potent antagonist failed to inhibit the increased afferent discharge post-CSD. Although activation of P2X2/3 by ATP may contribute to the afferent activation, the action of other algesic mediators that are co-released is likely more potent in driving the discharge of meningeal afferents post-CSD.

One methodological aspect of our study that could potentially affect data interpretation is the possible inhibitory effect of the CBX treatment on brain gap junctions and 11β-hydroxysteroid dehydrogenase (11β-HSD1). We found that topical dural CBX application did not affect the propagation of CSD, which is controlled by gap junctions (Theis et al., 2003). This data suggests that the concentration of CBX attained in the cortex following epidural administration, which is expected to be substantially lower due to poor transmeningeal diffusion (Zhao et al., 2017), was insufficient to inhibit cortical gap junctions. At present, we cannot exclude an effect mediated by increased brain levels of corticosterone due to 11β-HSD1 inhibition by CBX. Nonetheless, due to the restricted diffusion across the meninges discussed above, at the concentration used in the current study, CBX is unlikely to affect brain corticosterone levels (Bonvalet et al., 1990).

The absence of females is a limitation of this study, given the high prevalence of migraine in women. A recent study conducted in C57Bl/6 mice reported that P2X7 mediates muscle pain in males but not females (Hayashi et al., 2023). Of note, this mouse strain expresses a loss-of-function P2X7 variant that might have contributed to the sex differences observed. Another study in rats, however, implicated peripheral P2X7 activation in trigeminal pain in both sexes and further described estradiol-mediated increased P2X7 expression in females (Bereiter et al., 2022). Finally, non-genomic action of estrogen inhibits human P2X7 activation (Cario-Toumaniantz et al., 1998). Thus, the effects of sex and ovarian hormones on P2X7-mediated pain are likely complex. Additional studies in multiple species will be needed to determine whether the contribution of peripheral P2X7 to meningeal nociception is sexually-dimorphic and/or involves changing levels of estrogens.

Meningeal afferents are considered major players in migraine pain, yet our understanding of how they become engaged during a migraine attack is limited. We uncover a role for meningeal P2X7 signaling, involving Panx1 megachannels in mediating the mechanical sensitization of meningeal afferents in a model of migraine of aura. Therapeutic approaches and clinical trials of P2X7 blockers are currently under development for various inflammatory conditions (Rotondo et al., 2022) and may be further explored for migraine treatment.

## Acknowledgments

This work was supported by NIH grants: R01NS086830, R01NS115972, R01NS078263

